# Transcranial Magnetic Stimulation Reduces Non-Decision Time in Perceptual Decisions

**DOI:** 10.1101/2025.02.13.637859

**Authors:** Susanna Gordleeva, Vladimir Maksimenko, Usama Mohammad, Nikita Grigoriev, Andrey Savosenkov, Alexander Kuc, Semen Kurkin, Anna Udoratina, Vadim Grubov, Alexander Hramov

**Affiliations:** Neuromorphic computing center, Neimark University, Nizhny Novgorod, Russian Federation, 603081; Department of Neurotechnology, Lobachevsky State University, Nizhny Novgorod, Russian Federation, 603022; Faculty of Computer science and Engineering Innopolis University, 420500 Innopolis, The Republic of Tatarstan, Russia; Baltic Center for Neurotechnology and Artificial Intelligence, Immanuel Kant Baltic Federal University, Kaliningrad, Russian Federation, 236041; Centre for Cognition and Decision making, Institute for Cognitive Neuroscience, HSE University, Russian Federation, 101000

**Keywords:** Perceptual decision-making, perceptual learning, drift-diffusion model, non-decision time, transcranial magnetic stimulation, response time

## Abstract

Perceptual decision-making enables humans to process sensory information and translate it into goal-directed actions. A critical component of this process is response time, which declines with age due to the slowing of both non-decision processes and evidence accumulation. While perceptual learning has been shown to counteract these declines, it remains unclear whether non-decision processes can be accelerated through direct neural stimulation, bypassing the need for long training sessions. Here, we propose that Transcranial Magnetic Stimulation (TMS) can enhance perceptual decision-making by specifically reducing non-decision time. To test this hypothesis, we tracked response times during a perceptual learning task while recording brain activity. Using the Drift Diffusion Model, we quantified the effects of TMS on mental speed and non-decision time. Our findings reveal that perceptual learning decreases response time by simultaneously increasing mental speed and reducing non-decision time. Crucially, TMS application to brain regions associated with perceptual learning further shortened non-decision time without altering mental speed. These results demonstrate that TMS can selectively accelerate non-decision processes, offering a promising intervention for mitigating age-related cognitive slowing and enhancing decision efficiency.

**Significance Statement:** Perceptual decision-making is a fundamental cognitive process that slows with age due to declines in both non-decision processes and mental speed. While perceptual learning can mitigate these declines through practice, we propose that non-decision time can also be reduced by directly stimulating neural ensembles using Transcranial Magnetic Stimulation (TMS). Our study demonstrates that TMS applied to task-relevant brain areas decreases non-decision time, thereby accelerating response speed without affecting mental speed. These findings reveal a novel mechanism for enhancing perceptual decision-making efficiency and highlight TMS as a potential tool for counteracting age-related cognitive slowing.

## Introduction

Humans and animals process sensory information and translate it into actions - a process commonly referred to in the literature as perceptual decision-making (1-4). In visual perceptual decision-making, the time between the appearance of sensory information in visual field and the corresponding motor response is known as response time, an important characteristic of perceptual decisions. Response speed declines with age. Perceptual decision making involves two main components: non-decision processes and evidence accumulation. Non-decision processes associated with encoding sensory information and motor response formation start slowing down at the age of 20. The speed of evidence accumulation, which is known as mental speed, declines at the age of 60 (5).

The brain, however, can be trained to process information more efficiently and resist age-related slowing, a process known as perceptual learning (6). The core idea of perceptual learning is that performance on perceptual tasks improves with practice. In other words, repeated exposure to stimuli reduces the response time needed to process these stimuli in the future. Perceptual learning helps decrease response time by enhancing both components of perceptual decision-making: it reduces non-decision time (7) and increases mental speed (8). Mental speed, which is linked to higher-level cognitive processes, is known to be stimulus-specific; to increase mental speed, perceptual learning should involve interaction with the specific stimulus (9). In contrast, non-decision time involves more low-level processes such as attention allocation, stimulus encoding, memory access, and response execution, which remain consistent across different stimuli (7).

We propose that non-decision processes can also be reduced by directly stimulating neural ensembles without requiring interaction with the stimuli. We suggest using Transcranial Magnetic Stimulation (TMS) due to its safety and proven ability to induce sustained excitatory or inhibitory effects on neuronal ensembles (10, 11). We hypothesize that TMS will decrease non-decision time for two main reasons. First, TMS increases the probability of spiking activity, resulting in a higher speed of information processing in neural ensembles (12). Second, TMS promotes neural plasticity and facilitates functional reorganization within brain networks, which is crucial for perceptual learning (13). The ability of TMS to affect response time has been demonstrated (14), but the connection between TMS and non-decision time remains unexplored.

To test these hypotheses, we enrolled participants in a perceptual learning task while tracking response time. We recorded brain activity and identified areas where the power of brain signals correlates with perceptual learning. We applied TMS to these areas before the experiment and test whether the response time changed. Finally, we used the Drift Diffusion Model to estimate the effect of TMS on mental speed and non-decision time. We show that perceptual learning reduces response time, increases mental speed, and decreases non-decision time. These processes are associated with increased spectral power of brain signals over the frontal cortical areas. Application of TMS to these areas reduces response time as well as non-decision time but does not affect mental speed. Our results confirm the potential of TMS to decrease non-decision time, thereby contributing to an overall increase in the speed of perceptual decision-making.

## Results

### Perceptual Learning and DDM Parameters

Participants (n = 30) took part in the experiment where they repetitively responded to visual stimuli presented with brief inter-stimulus intervals, while we collected their response time (RT) and accuracy (see Methods). We observed high accuracy (M = 0.873, SD = 0.113), evidencing that they made informed decisions based on sensory evidence rather than randomly guessing. To confirm perceptual learning, we were interested if the behavior of participants changed with the time spent in the experiment; therefore, we separated all stimuli into four blocks according to their presentation times (see Methods).

To explain changes in behavior, the drift-diffusion model (DDM) was fitted to the choice and response time data for each block for each participant. The DDM assumes that when looking at the stimulus with two possible interpretations, participants accumulate evidence in favor of each interpretation. When the evidence accumulated for one interpretation reaches the decision threshold, this interpretation is chosen by a participant (See Fig. 1, A). It is assumed that participants start accumulating evidence sometime after stimulus presentation. The time before evidence accumulation is known as non-decision time (τ_0_) and includes some basic processes associated with attention allocation and information encoding. Then, it is assumed that evidence is accumulated at a constant speed, known as drift rate (μ). Since RT varies across trials, the model adds a Gaussian noise term to the drift rate, known as diffusion term with dispersion (σ). The parameters, including drift rate, non-decision time, and decision threshold, are estimated from the data to ensure that the drift-diffusion model can fit and predict response time and accuracy.

**Figure 1.**
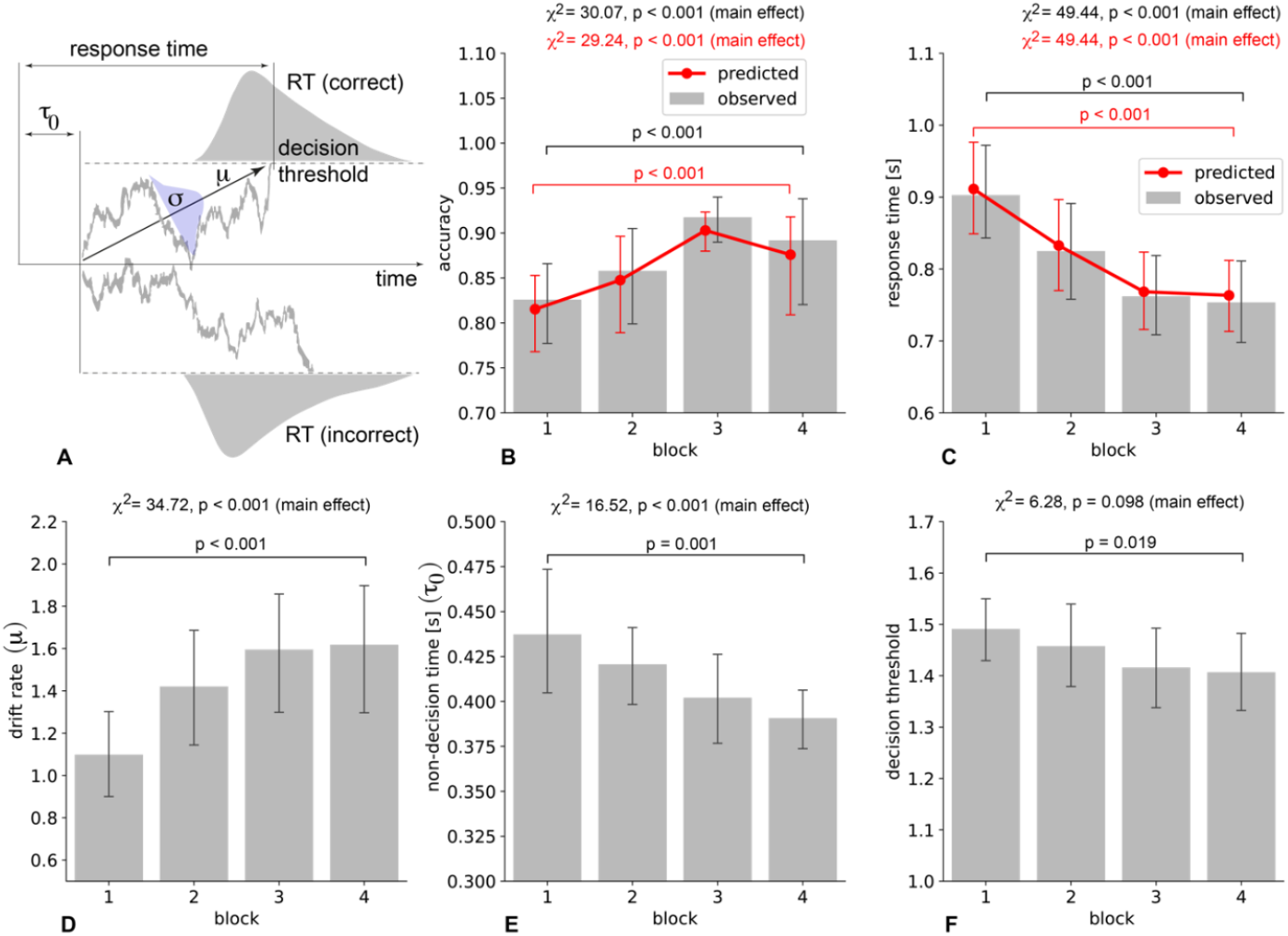
Behavioral data and DDM parameters. (A) Schematic illustration of the evidence accumulation process in DDM and its parameters including non-decision time (τ_0_), drift rate (μ), noise term (σ), and decision threshold. RT distributions illustrate response times observed in the experiment. (B) Accuracy (observed in experiment and predicted by DDM) for all blocks. (C) Response time (observed and predicted) for all blocks. Estimated DDM parameters, drift rate (D), non-decision time (E), and decision threshold for all blocks. Data are shown as group mean and 95%CI based on 1000 bootstrap iterations. Main effect of block is estimated using Fieldman test and difference between block 1 and block 4 is estimated based on Wilcoxon test. All p-values are uncorrected.

First, we observed that behavioral responses of participants changed between blocks. The mean accuracy showed a tendency to increase, and this tendency was significant: χ^2^(3) = 30.07, p <0.001 (Fig. 1, B). The post-hoc analysis revealed that accuracy in block 1 (M = 0.825, SD = 0.126) was significantly lower than in block 4 (M = 0.892, SD = 0.173): z = −3.835, p <0.001. Similarly, RT showed a tendency to decrease, and this tendency was also significant: χ^2^(3) = 49.44, p <0.001 (Fig. 1, C). Post-hoc analysis revealed that RT in block 1 (M = 0.902 s, SD = 0.189) was significantly higher than in block 4 (M = 0.753 s, SD = 0.154): z = −4.658, p <0.001.

Second, we observed that the DDM was able to fit and predict response time and choices of participants. To assess predictive performance, we estimated correlation between the observed and predicted RTs and accuracies and found a strong positive correlation between observed and predicted RTs: r(88) = 0.994, p < 0.001 95%CI [0.99, 1.0], as well as between predicted and observed accuracies: r(88) = 0.996, p < 0.001 95%CI [0.99, 1.0]. Similar to the observed accuracy, predicted values showed a significant tendency to increase: χ^2^(3) = 29.24, p <0.001, so that predicted accuracy in block 1 (M = 0.815, SD = 0.121) was lower than in block 4 (M = 0.875, SD = 0.163): z = −3.77, p < 0.001 (Fig. 1, B). Similar to the observed RT, predicted values showed a significant tendency to decrease: χ2(3) = 49.44, p <0.001, so that predicted RT in block 1 (M = 0.911 s, SD = 0.183) was higher than in block 4 (M = 0.763, SD = 0.145): z = −4.573, p < 0.001 (Fig. 1, C).

Finally, we found that DDM parameters change between blocks. Drift rate shows tendency to increase, and this tendency was significant: χ^2^(3) = 29.24, p <0.001 (Fig. 1, D). Post-hoc analysis revealed that drift rate in block 1 (M = 1.098, SD = 0.571) was significantly lower than in block 4 (M = 1.618, SD = 0.855): z = −4.123, p < 0.001. Non-decision time showed tendency to decrease, and this tendency was also significant: χ^2^(3) = 29.24, p <0.001 (Fig. 1, E). Post-hoc analysis revealed that non-decision time in block 1 (M = 0.437, SD = 0.096) was significantly higher than in block 4 (M = 0.391, SD = 0.051): z = −3.013, p = 0.001. Finally, the decision threshold did not change across blocks: χ^2^(3) = 6.28, p = 0.098 (Fig. 1, F).

Together, these results suggest that participants improve their responses with the time spent in the experiment, showing increasing accuracy in stimulus recognition and decreasing response time. These changes were predicted by the DDM and were associated with changes in DDM parameters, including growing drift rate and decreasing non-decision time.

### Perceptual Learning and Brain Response

To reveal brain activity associated with the observed changes in behavior, we compared the spectral power of EEG signals in the frequency range of 1 – 40 Hz time-locked to the stimulus onset (Fig. 2, A) and to the button pressing (Fig. 2, B). We identified three time-intervals of interest (TOIs) and compared spectral power across blocks at each TOI using permutation statistics over EEG sensors and frequencies (see Methods). For TOI1, we revealed a cluster of EEG sensors (Cz, T8, FC1, FC2, FC6, FT10, F3, Fz, F4, F8, Fp1, Fp2), where spectral power in the frequency band of 5 – 12.25 Hz significantly differed across blocks (p = 0.0009). In TOI2, we revealed a cluster of EEG sensors (CP2, C3, Cz, C4, T8, FC1, FC2, FC6, FT10, F7, F3, Fz, F4, F8, Fp1, Fp2), where spectral power in the frequency band of 8.25 – 12 Hz significantly differed across blocks (p = 0.0029). Finally, in TOI3 we did not find a significant change in EEG spectral power across blocks. In TOI1 and TOI2, spectral power increased from block 1 to block4 (Fig. 2, D).

**Figure 2.**
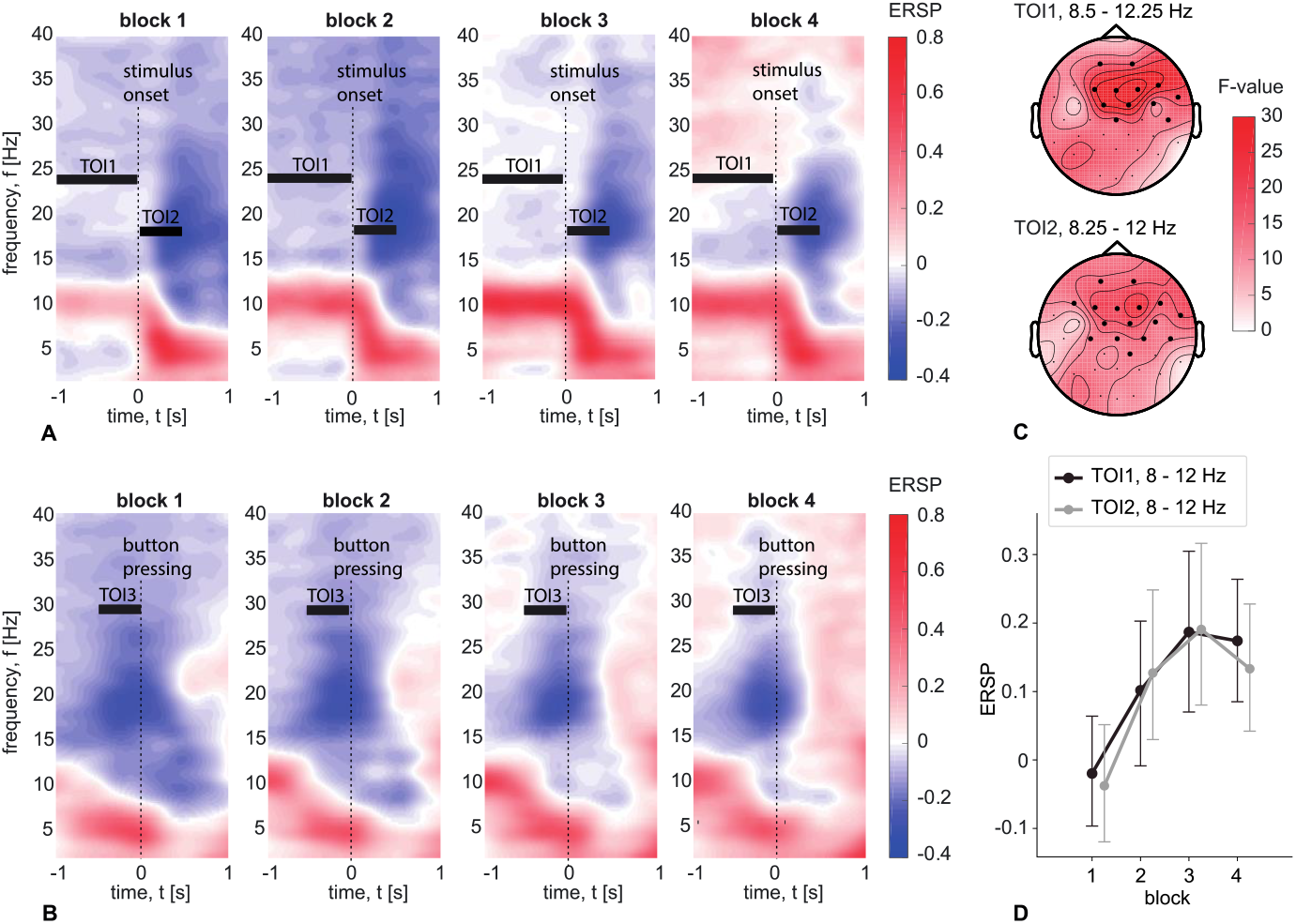
Changes in brain response with time on task at the sensor level. Time-frequency distribution of the spectral power of EEG signals is shown for the time intervals locked to stimulus onset (A) and button press (B) across four blocks. Data are presented as the grand average across stimuli, electrodes, and participants. Time intervals of interest (TOI1, TOI2, and TOI3) are indicated by black rectangles. Topographic plots (C) depict the distribution of F-statistics resulting from the comparison of 8–12 Hz spectral power at TOI1 and TOI2 between blocks. EEG sensors where the statistics surpass the significance threshold are highlighted. The point plot (D) illustrates 8–12 Hz spectral power at TOI1 and TOI2, averaged across these sensors for four blocks. Data are shown as the mean across participants with 95% confidence intervals.

The results obtained for TOI1 and TOI2 show an overlap in frequency band and EEG sensors. Therefore, we combined TOI1 and TOI2 and calculated the distribution of EEG power in the frequency band of 8.5 – 12 Hz across the voxels in the source space. Comparing the EEG power in the source space between blocks, we localized a cluster of voxels where activation changed significantly between blocks (see Fig. 3, A). Finally, we estimated whether the changing EEG power in these zones is associated with the observed behavioral changes and model parameters.

**Figure 3.**
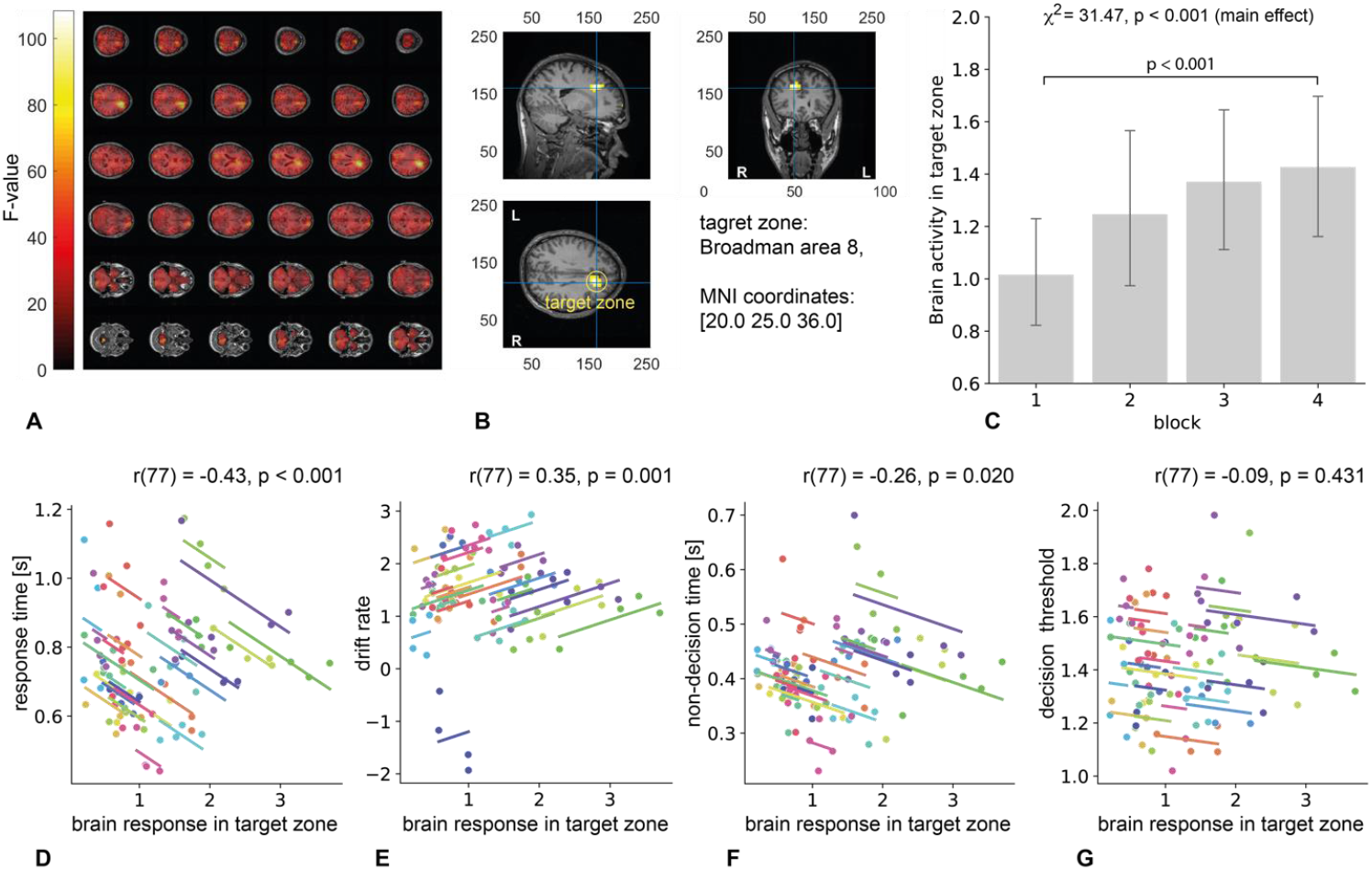
Brain activity and its association with behavior and DDM parameters. (A) Distribution of the F-statistics (based on one-way ANOVA) across voxels reflecting difference in EEG power between blocks. (B) The cluster of voxels (target zone) where the difference in EEG power between blocks is significant (p < 0.001 corrected with permutation framework). (C) Distribution of EEG power in the target zone across blocks. For each block, data are shown as group mean and 95%CI based on 1000 bootstrap iterations. Main effect of block is estimated using Fieldman test and difference between block 1 and block 4 is estimated based on Wilcoxon test. All p-values are uncorrected. (D-E) Correlation between the EEG power in target zone, response time, and DDM parameters. Data are shown as a set of participant-level regression lines, having the same slope and scatterplot showing the participant-level observations (different colors correspond to different participants).

We observed only one cluster of voxels showing a significant change in EEG power between blocks, which corresponded to the highest values of the F-statistic (Fig. 3, A) When applying Automated Anatomical Labeling (AAL), we found that this cluster corresponded to the Broadman area 8 (Fig. 3, B). The EEG power averaged over the voxels in this area showed a tendency to increase across blocks: χ^2^(3) = 31.47, p <0.001, so that the power in block 1 (M = 1.012, SD = 0.581) was significantly lower than in block 4 (M = 1.415, SD = 0.777): z = −4.051, p < 0.001 (Fig. 1. C).

We observed a moderate negative correlation between EEG power in the right anterior cingulate cortex and RT: r(77) = −0.431, p < 0.001, CI95%[-0.6, −0.23] (Fig. 3, D); a moderate positive correlation between EEG power and drift rate: r(77) = 0.353, p = 0.001, CI95%[0.14, 0.53] (Fig. 3, E), and a weak negative correlation between EEG power and non-decision time: r(77) = −0.261, p = 0.021, CI95%[-0.46, −0.04] (Fig. 3, F). No correlation was observed between the EEG power in the right anterior cingulate cortex and decision threshold: r(77) = −0.089, p = 0.431, CI95%[-0.3, 0.13] (Fig. 3, G).

Together, these results show that activity of the right anterior cingulate cortex at TOI1 and TOI2 grows with the time spent on a task and that this growth is associated with decreasing RT, growing drift rate, and decreasing non-decision time.

### Effect of TMS on Behavior and Model Parameters

To test the effect of TMS, we randomly divided respondents into a Control group (n = 15) and a Treatment group (n = 15). Six months after the experiment described above (which we refer to as the baseline session), respondents participated in a similar experiment (which we refer to as the stimulation session). The Control group received SHAM stimulation while the Experimental Group received repetitive TMS stimulation applied to the Broadman area 8 before participating in the experiment (Fig. 4, A). First, we estimated whether the RT and accuracy of respondents changed between baseline and stimulation sessions in both the Control and Treatment groups. Second, we fit the DDM to the experimental data and tested whether the predicted RT and accuracies differed between the baseline and stimulation sessions in both groups. Finally, we compared DDM parameters between the baseline and stimulation sessions.

**Figure 4.**
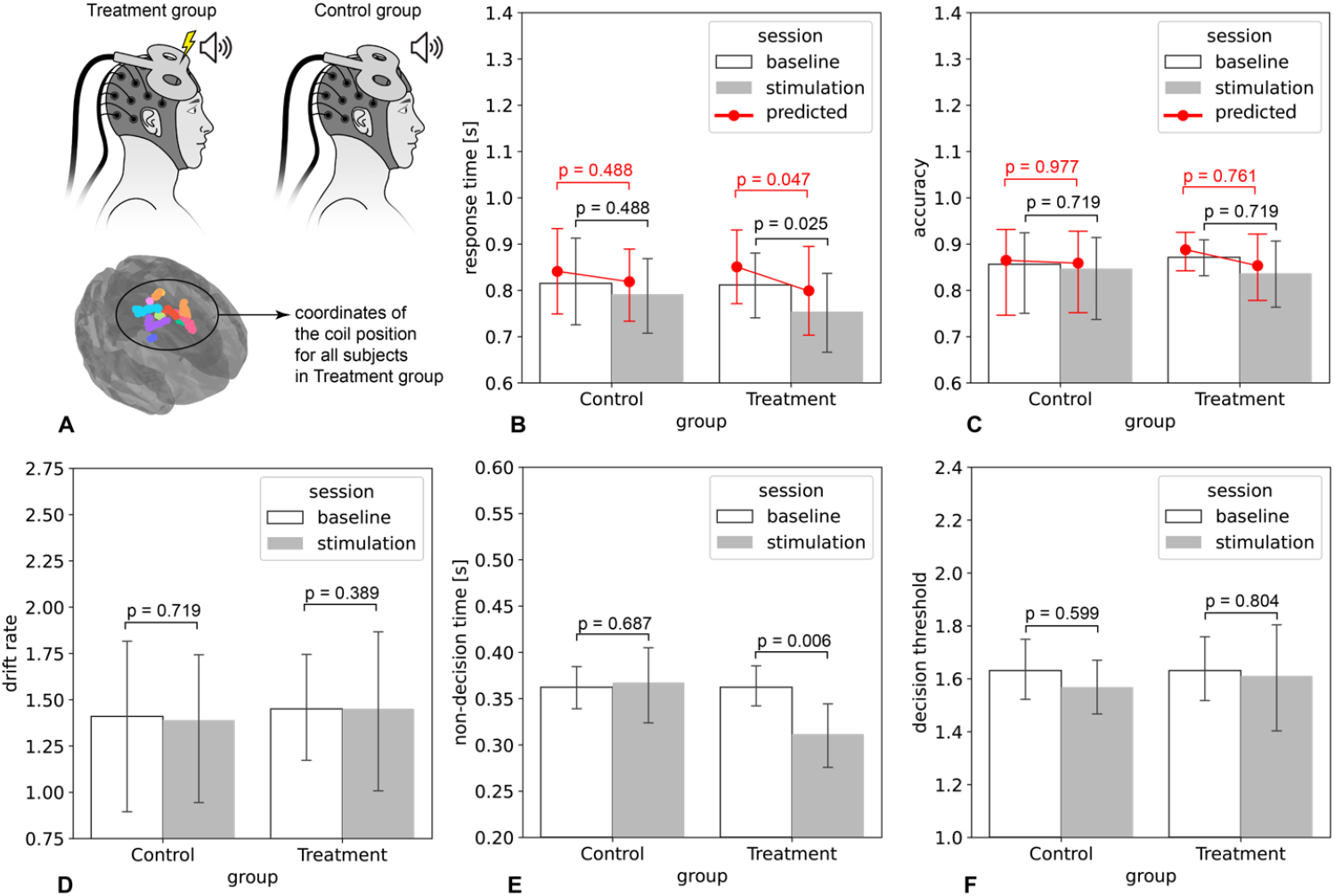
Effect of TMS on behavior and DDM parameters. (A) Schematic illustration of TMS stimulation in Treatment group and SHAM stimulation Control group as well as coordinates of the TMS coil in the stimulation session of treatment group (different colors correspond to different participants). (B) Comparison of observed and predicted response times between baseline and stimulation sessions in both groups. (C) Comparison of observed and predicted accuracies between baseline and stimulation sessions in both groups. (D-E) Comparison of estimated model parameters between baseline and stimulation sessions in both groups (individual models are estimated for each participant). Data are shown as group mean and 95%CI based on 1000 bootstrap iterations. Significance is estimated based on Wilcoxon test. All p-values are uncorrected.

In the Control group, there was no difference in RT between the baseline (M = 0.815 s, SD = 0.189) and stimulation (M = 0.792 s, SD = 0.163) sessions: z = −0.738, p = 0.488. In the Treatment group, RT was lower in stimulation (M = 0.754 s, SD = 0.176) compared to the baseline (M = 0.811 s, SD = 0.146) session: z = −2.215, p = 0.025 (Fig. 4, B). We further estimated Bayesian statistics for this case and obtained B10 = 5.55 indicating moderate to strong evidence in favor of the alternative hypothesis (the difference in RT between sessions) over the null hypothesis (RT remains same across sessions). Accuracy in the Control group did not differ between the baseline (M = 0.872, SD = 0.088) and stimulation (M = 0.837, SD = 0.147) sessions: z = −0.397, p = 0.719. In the Treatment group, accuracy also did not differ between the baseline (M = 0.856, SD = 0.196) and stimulation (M = 0.847, SD = 0.189) sessions: z = −0.397, p =0.719 (Fig. 4, C).

The DDM predicted similar tendencies for RT and accuracy. In the Control group, there was no difference in the predicted RT between the baseline (M = 0.841 s, SD = 0.186) and stimulation (M = 0.818s, SD = 0.155) sessions: z = −0.738, p = 0.488. In the Treatment group, predicted RT was higher in baseline (M = 0.815 s, SD = 0.189) than in stimulation (M = 0.792 s, SD = 0.163) sessions: z = −1.987, p = 0.047 (Fig. 4, B). Predicted accuracy in the Control group did not differ between the baseline (M = 0.887, SD = 0.083) and stimulation (M = 0.853, SD = 0.149) sessions: z = −0.341, p = 0.761. In the Treatment group, accuracy also did not differ between the baseline (M = 0.865, SD = 0.206) and stimulation (M = 0.858, SD = 0.195) sessions: z = −0.057, p =0.977 (Fig. 4, C). Considering DDM parameters, we observed that drift rate in the Control group did not differ between the baseline (M = 1.410, SD = 0.899) and stimulation (M = 1.390, SD = 0.802) sessions: z =-0.397, p = 0.719. In the Experimental group, drift rate also did not differ between the baseline (M = 1.451, SD = 0.594) and stimulation (M = 1.452, SD = 0.898) sessions: z =-0.908, p = 0.389. In the Control group, there was no difference in non-decision time between the baseline (M = 0.362 s, SD = 0.047) and stimulation (M = 0.367 s, SD = 0.078) sessions: z = −0.454, p = 0.687. At the same time, non-decision time was lower in the stimulation session (M = 0.311 s, SD = 0.071) compared to the baseline session (M = 0.363 s, SD = 0.071) in the Treatment group: z = −2.612, p = 0.006 (Fig. 4, D). The decision threshold in the Control group did not differ between the baseline (M = 1.631, SD = 0.241) and stimulation (M = 1.569, SD = 0.213) sessions: z = −0.567, p = 0.599. In the Treatment group, the decision threshold also did not differ between the baseline (M = 1.615, SD = 0.204) and stimulation sessions (M = 1.611, SD = 0.404): z = −0.283, p =0.804 (Fig. 4, E) Together, these results suggest that repetitive TMS applied to the Broadman area 8 before the experiment decreases RT and reduces non-decision time, while accuracy, drift rate, and decision threshold do not change.

## Discussion

The analysis of behavioral data indicates that, during a perceptual decision-making task involving repeated presentations of similar visual stimuli, participants demonstrate progressively shorter response times and increased response accuracy. Drift-diffusion modeling reveals an increase in drift rate and a reduction in non-decision time, while the decision threshold remains constant. Drift rate, which reflects evidence accumulation, is determined by interactions with the stimuli and is unaffected by accompanying sensory-motor processes (15-17). Non-decision time, in contrast, represents processes unrelated to decision-making, such as information encoding and motor response formation (18, 19). The decision threshold, representing the amount of information required to make a decision, provides insight into participants’ task engagement. For instance, a reduction in the threshold is typically associated with a shift from accuracy-focused to speed-focused decision-making (20-22). These findings suggest that participants exhibit perceptual learning, characterized by enhancements in both low-level processes, such as stimulus encoding and motor response formation, and high-level processes associated with evidence accumulation during stimulus interaction. Notably, the lack of change in the decision threshold indicates that perceptual learning does not alter the speed-accuracy tradeoff or participants’ task-related attitudes. These results are consistent with observations reported by Petrov, Van Horn, and Ratcliff (7).

The analysis of brain activity demonstrates a progressive increase in spectral power within the 8– 12 Hz frequency band as the experiment proceeds. This effect is consistently observed during the time intervals immediately preceding and following stimulus onset, localized to the same EEG sensor region. These findings suggest that neural activity in this area may be elicited by the preceding stimulus and persist throughout the inter-stimulus interval, potentially facilitating a more rapid and efficient activation in response to subsequent stimuli. The observed increase in EEG spectral power was not evident during the time interval preceding the response to the stimulus. Prior research indicates that, in perceptual decision-making tasks, cognitive processes associated with sensory processing and decision-making are temporally dissociated. Sensory processing predominantly occurs within the first 300 ms following stimulus onset, whereas decision-making processes become more prominent closer to the response onset (23-25). Accordingly, we propose that the observed effect reflects the activity of neural ensembles engaged in cognitive processes during the sensory processing phase rather than during the decision-making stage.

Source-space analysis of brain activity indicates that the observed increase in spectral power is localized to the right side of Broadman area 8, which according to literature is as part of the right dorsolateral prefrontal cortex (rDLPFC) (26-30). Previous research has established the rDLPFC as a critical region involved in various components of visual information processing, including cognitive control functions (31, 32), adjustment of decision-making strategies, cognitive flexibility (33-35), and filtering of decision-relevant information (36; 37). Of particular interest is the relationship between increased spectral power in the 8–12 Hz frequency band and improved task performance. Neural activity within this band is conventionally associated with relaxation and classified as task-negative, where task engagement typically corresponds to decreased spectral power in this band (38, 39). However, there is evidence that 8–12 Hz activity can also serve a task-positive role, characterized by an increase in activity linked to enhanced performance. This task-positive activity is hypothesized to facilitate the inhibition of irrelevant information, thereby enhancing the extraction and processing of decision-relevant features from sensory input (40-42). Notably, elevated alpha-band activity in the prefrontal cortex has been previously associated with the selection of decision-relevant information (43). Considering these findings, we propose that perceptual learning enhances the capacity of neural ensembles to filter decision-relevant information during sensory processing. This improvement is reflected in the observed increase in alpha-band activity within the prefrontal cortex.

Transcranial magnetic stimulation (TMS) applied to the right dorsolateral prefrontal cortex (rDLPFC) was found to reduce response time, specifically by decreasing non-decision time, as estimated through the drift-diffusion model. Other parameters, however, remained unaffected. This outcome supports our hypothesis that direct brain stimulation via TMS can effectively decrease non-decision time. As expected, the drift rate, representing the more complex processes underlying the interaction between the respondent and the stimuli, was not influenced by TMS, indicating that this parameter is less susceptible to modulation through direct cortical stimulation. Non-decision time typically encompasses processes related to the encoding of sensory information and the formation of a motor response following the decision-making phase. Brain activity analysis indicated that changes were observed in the interval following the onset of the stimulus, rather than before the motor response. These findings suggest that TMS specifically affects the encoding component of non-decision time, which involves processing visual information about the stimulus prior to the accumulation of evidence. This insight implies that TMS could potentially reduce response time for a wide range of stimuli, as the early stage encoding processes following stimulus onset are likely to be independent of the specific characteristics of the stimulus. Moreover, the reduction in response time was shown in younger participants, for whom the decline in response time is generally minimal (5). It is anticipated that TMS will exert a more substantial effect in older adults, who exhibit greater slowing of response times and extended non-decision processes (44). This suggests that TMS may serve as a valuable intervention for mitigating age-related cognitive slowing, offering a potential means of compensating for the decline in response speed and non-decision processing typically observed with aging.

## Materials and Methods

### Participants

Thirty healthy subjects (16 females and 14 males) aged 18 to 33 years (M = 21.5, SD = 2.8), were recruited from the students of Lobachevsky State University of Nizhny Novgorod. This sample size falls within the range used in experiments with transcranial magnetic stimulation (45-53). The experiment’s methodology was developed in accordance with the Declaration of Helsinki and approved by the Local Ethical Committee of Lobachevsky State University of Nizhny Novgorod (protocol No. 2, dated March 19, 2021). All subjects underwent an interview to identify any contraindications to Transcranial Magnetic Stimulation (TMS) and were informed about the potential side effects. All volunteers provided written informed consent.

### Visual stimuli and Experimental Task

We employed a perceptual decision-making task involving the perception of ambiguous visual stimuli, specifically Necker cubes. A Necker cube is a 2-D drawing of a cube with visible internal edges, which observers interpret as a 3-D object oriented either to the left or to the right. By adjusting the contrast of the inner edges, one can make either the left-oriented or right-oriented projection dominate. When all internal edges are equally visible, the cube becomes ambiguous; that is, its projection is not objectively identifiable. In our study, ambiguous cubes were excluded to allow participants to make informed decisions regarding the stimulus orientation. We utilized eight Necker cubes (Fig. 5, A), distinguished by the parameter *a* ∈ [0,1], which reflects the normalized edge luminance in a grayscale palette. Consequently, the brightness of the inner edges forming the left-lower and right-upper squares on the 2D image was calculated as *l=1-a*, and *r=a*, respectively.

**Figure 5.**
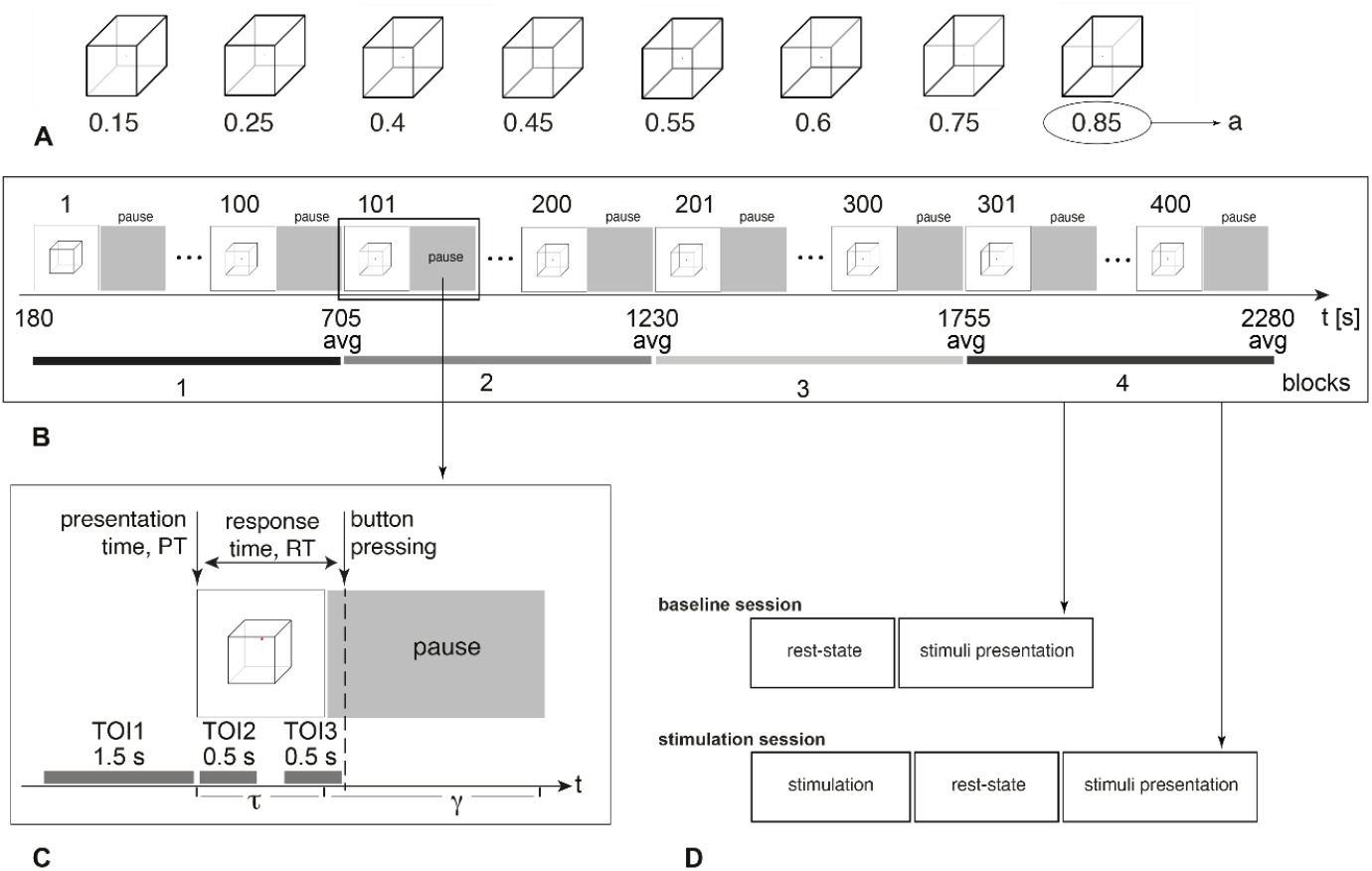
Experiment Overview. (A) Visual stimuli set: Necker cubes used in the experiment, with parameter *a* controlling the contrast of the inner edges. (B) Experimental session structure: 400 stimuli were presented in random order, divided into 4 blocks, each containing 100 stimuli. (C) Presentation structure: presentation time varied from 1 to 1.5 seconds, while the pause between presentations ranged from 5 to 5 seconds; response time (RT) is defined as the duration between stimulus onset and button press. (D) Overview of the baseline and stimulation session structures: Baseline and Stimulation sessions had the same structure excepting stimulation applied before the rest-state.

Participants were comfortably seated in a chair, holding a two-button input device connected to the amplifier with both hands. They were instructed to press either the left or right key when recognizing the left or the right stimulus orientation. Necker cube images of 25.6 cm were displayed on a 27-inch LCD screen (with a resolution of 1920×1080 pixels and a 60 Hz refresh rate) located at a distance of 2 meters from the participant. Each stimulus appeared on the screen for a short time interval, randomly chosen from the range of1-1.5 s. Between the stimuli, we demonstrated an abstract image for 3-5 s. The timing of stimuli presentations and the EEG streams were synchronized using a photodiode connected to the amplifier. During experimental sessions, the cubes with predefined ambiguity were randomly demonstrated 400 times. The experiment lasted around 38 min (Fig. 5, B, C).

### Transcranial Magnetic Stimulation

For stimulation, we used a TMS Neuro-MS/D Advanced Therapeutic system (Neurosoft, Russia) with an AFEC-02-100-C cooled angulated figure-of-eight coil (100 mm). To navigate the coil, we used an infrared marker set to the target zone. The coil was placed and fixated over the activation zone, and the handle was angled 45° to the longitudinal cerebral fissure. We applied excitatory stimulation with the following parameters: 1800 stimuli, 10 Hz, 3 minutes. For SHAM stimulation, all parameters were the same, but the coil was placed on its wing (90° relative to the head) to stimulate away from the head. Stimulation started with the calibration procedure. We used the TMS navigator’s system (Localite, Germany) to generate the 3D image of the participant’s brain and mark the target zone on it. To find the individual motor threshold, we performed a set of 10 stimulations of the motor cortex with different power levels. The power level at which the evoked motor potentials arose in 5/10 stimulations was taken as an individual motor threshold. Further simulations were set on 120% of individual motor threshold power.

### Experiments

Participants were divided into two equal groups: Control and Treatment. In both groups, each participant took part in two sessions, Baseline and Stimulation, with a 2–3-month break between sessions (Fig. 5, D). In the Baseline session, participants underwent the MFI test to estimate fatigue levels. Then, they underwent a 5-minute recording of the rest-state EEG. Finally, they participated in a stimuli classification procedure described above. In the Control session, participants completed the MFI test, then they were subjected to TMS stimulation followed by a 3-minute recording of the rest-state EEG and the same stimuli classification procedure. The Control group underwent SHAM stimulation, while the Experimental group received excitatory TMS stimulation to the target zone.

### Brain Activity Recording and Processing

We registered electroencephalograms (EEG) using a 48-channel NVX-52 amplifier (MKS, Zelenograd, Russia). EEG signals were recorded from 32 standard Ag/AgCl electrodes placed according to the 10-20 layout. The earlobe electrodes were used as a reference. The ground electrode was placed on the forehead. Impedance was kept below 10 KΩ. EEG was digitized with a sampling rate of 1000 Hz. A band-pass FIR filter filtered the raw EEG signals with cut-off points at 1 Hz (HP) and 100 Hz (LP) and with a 50-Hz notch filter. Eye-blinking and heartbeat artifacts were removed by Independent Component Analysis using EEGLAB software.

We calculated wavelet power (WP) of EEG signals in the frequency band of 1−40 Hz using the Morlet wavelet. The number of cycles, denoted as n, depended on the signal frequency f as n=f. Stimuli with erroneous responses and fragments containing artifacts with WP exceeding ±3 standard deviations of the mean WP were excluded from the analysis. For the remaining data, we analyzed WP in three time-intervals of interest (TOI): TOI1 was a 2-second interval preceding stimulus onset, TOI2 was a 0.5-second interval following stimulus onset, and TOI3 was a 0.5-second interval preceding button pressing (see Fig. 5, C). The WP in these TOIs were normalized using WP obtained in the rest-state as (WP−WPrest)/WPrest. All calculations were performed using the Fieldtrip toolbox in MATLAB.

To estimate EEG power in the source space (referred to as the source power, SP), we re-referenced EEG signals to the common average and subtracted the mean. We solved the inverse problem using Dynamical Imaging of Coherent Sources (54). The 3D head model was created using the ‘Colin27’ brain MRI averaged template (55) and the boundary element method (56). As a result, we considered SP in the source space consisting of 11,929 grid points (voxels) inside the brain. Finally, we normalized the obtained SP estimates at each voxel to the SP estimated for the resting state. The automated anatomical labeling brain atlas (57) was used to map the location of voxels to anatomical brain regions.

### Drift-Diffusion Model Estimation

We estimated three parameters of the Drift Diffusion Model (DDM) from the data: drift rate, non-decision time, and decision threshold. Parameter estimation was conducted using the Python package rlssm (58), which implements Bayesian parameter estimation. Specifically, the package employs the No-U-Turn Sampler (59), a variant of Hamiltonian Monte Carlo (60), to perform the Bayesian estimation.

Individual-level models were estimated for each participant, rather than employing a hierarchical modeling approach. To evaluate the model’s fitting performance, we utilized the Watanabe-Akaike Information Criterion (61). WAIC is a generalized extension of the Akaike Information Criterion (62), designed to accommodate non-normal posterior distributions. Lower WAIC values indicate superior model fit in all cases.

### Hypothesis testing plan

First, we analyzed the response time (RT) data collected from 30 participants in the Baseline session. The experimental session was divided into 4 blocks, each including 100 subsequently presented stimuli (see Fig. 5, B). For each participant, we averaged the RT across the stimuli presented in each block. We then compared the obtained values between blocks using the nonparametric Friedman test, with a significance level set to 0.05. We conducted post-hoc pairwise comparisons of RT between blocks using nonparametric Wilcoxon test. To correct for multiple comparisons, the significance level was adjusted to 0.015.

Second, we fit the DDM to the response time and choice data of each participant in each block. We compared the response times and accuracies predicted by the DDM as well as estimated drift rate, non-decision time, and decision boundary across the 4 blocks using the Friedman test, with a significance level set at 0.05. We conducted post-hoc pairwise comparisons of each parameter between blocks using the nonparametric Wilcoxon test. To correct for multiple comparisons, the significance level was adjusted to 0.015.

Third, we analyzed EEG data collected from 30 participants during the Baseline session. For each block, we averaged wavelet power (WP) at three time-intervals of interest (TOIs) across the stimuli in each block (Fig. 5, C). After that, we compared WP at each TOI across blocks using F-test with the significance level set to 0.01. WP was distributed across sensors and frequencies, resulting in a large number of multiple comparisons. For correction, we employed permutation framework (63).

The significance level for the permutation statistics was set at 0.05. The number of permutations was set to 5000. These calculations were performed using the Fieldtrip toolbox in MATLAB.

This approach identified the subspaces of EEG sensors and frequencies (referred to as clusters) where WP differed significantly between blocks. For the post-hoc analysis, we averaged WP across channels and frequencies in each cluster and compared these values between blocks using the nonparametric Wilcoxon test. To correct for multiple comparisons, the significance level was adjusted to 0.015. This analysis was conducted in SPSS Statistics. Finally, we fixed the frequency bands of all clusters and performed a source reconstruction procedure to determine the SP distribution across brain tissue. These distributions were obtained for each block and compared between blocks using F-test with permutation framework. As a result, we identified groups of voxels where SP changed significantly between blocks (referred to as regions of interest, ROI). To associate changes of the SP in these ROIs with the changes in RT and DDM parameters, we used correlation analysis with repeated measures (64). Significance level was set to 0.01.

Fourth, we assessed whether the behavioral performance varied between the Baseline and Stimulation sessions for both the Control and Treatment groups. For each session, the mean RT was determined by averaging RTs across all stimuli. The correctness rate (CR) was calculated as the number of stimuli for which respondents correctly identified the orientation, divided by the total number of stimuli. Each of these metrics was compared between the Baseline and Stimulation sessions using the nonparametric Wilcoxon test with the significance level set to 0.05.

Fifth, we assessed whether the DDM parameters varied between the Baseline and Stimulation sessions for both the Control and Treatment groups. For each participant we fit DDM to the RT and choice data and evaluated the goodness of fit by computing spearman correlation between observed and predicted values. Then, each parameter was compared between the Baseline and Stimulation sessions using the nonparametric Wilcoxon test with the significance level set to 0.05.

## Author Contributions

S.G., V.M. and A.H. designed research; N.G., A.S. and A.U. collected the data; S.G. guided data acquisition; V.M., S.K., A.C., U.M., V.G. analyzed the data, with supervision from A.H.; V.M., S.K., S.G. interpreted the data; S.G. and V.M. wrote the manuscript.

## Acknowledgments

This work was supported by the Ministry of Science and Higher Education of the Russian Federation according to research project No. FSWR-2025-0009.

